# Cause and effect in the natural selection of the population ecological life histories of mammals

**DOI:** 10.1101/2024.03.23.586396

**Authors:** Lars Witting

## Abstract

The inter-specific life history and ecological variation of mammals is often explained as allometric consequences of physiological adaptations to unexplained body mass variation. But these hypotheses are unnecessary because the allometric scaling is explained already by the natural selection that explains the variation in mass. I decompose the population ecological life histories of 4,936 species of mammals to show how the selection of mass accounts for the life history and population ecological variation in mammals. This shows that 55% of the within order variance, and 91% of the between order differences, in the body mass, demography, and population ecological traits are reconciled by the response of population dynamic feedback selection to variation in net energy, mortality, and intra-specific interactive competition.

## 1 Introduction

Having body masses that span seven orders of magnitude, the life histories of mammals are often seen as consequences of physiological adaptations to size. This hypothesis was initially developed for metabolism by Rubner (1883), and extended to include life history traits like longevity and reproduction, and population ecological traits like population density, growth, and home range (e.g. Fenchel 1974; Damuth 1981; Peters 1983; Calder 1984; Brown et al. 2004).

Allometric correlations—where traits are described as linear functions of mass on double logarithmic scale—support the view, with an increasing number of physiological scaling hypotheses (e.g. Rubner 1883; Davison 1955; McMahon 1973; Blum 1977; Sibly and Calow 1986; West et al. 1997, 1999a,b; Banavar et al. 1999; Dodds et al. 2001; Dreyer and Puzio 2001; Rau 2002; Fujiwara 2003; Demetrius 2003; Santillán 2003; Ginzburg and Damuth 2008; Glazier 2010) generating a lack of consensus on the reason for metabolic scaling, and allometric correlations in general (e.g. Glazier 2005, 2010; West and Brown 2005; Witting 2008; White and Kearney 2013).

A main reason for our lack of ability to agree on a mechanism is the contingent paradigm, where biological evolution cannot be predicted forward, but can only be understood backwards after time’s actual unfolding (Mayr 1988; Salthe 1989; Gould 2002). This principle underlies traditional evolutionary thinking (including work by Lack 1947; Charlesworth 1980, 1994; Harvey and Pagel 1991; Roff 1992, 2002; Stearns 1992; Charnov 1993; Sibly and Brown 2007) that studies natural selection top-down backwards by analysing the fitness effects, trade-offs, and constraints in the evolved species of today.

The contingent approach underlies all physiological scaling hypotheses at some level, explaining metabolism as a dependent trait that follows from physiological traits (like vascular transportation networks; West et al. 1997) that adapt, or scale, to independent variation in body mass. But this method is problematic because it does not reconcile the natural selection of the dependent traits with the natural selection of the independent traits (Witting 1997, 2008). Contingent studies do not show that the independent traits (like mass & transportation networks) are selected by a deeper more fundamental natural selection than the selection that explains the dependent traits (like metabolism & transportation networks). It is therefore unresolved why the transportation network it not just selected to match a mass and metabolism that are selected by other means (Witting 1998).

Contingency furthermore allows for almost unlimited freedom; where we can choose more or less arbitrary among traits and trade-offs to explain a trait like metabolism, generating a potential multitude of natural selection hypotheses, with no theoretical guidance to separate realistic and unrealistic models from one another. It is therefore not surprising that each research team has its own favourite metabolic scaling hypothesis that has not been shown to be consistent with the natural selection of metabolism and mass.

A way forward is to restrict accepted hypotheses to models that show that they are self-sustained from a natural selection point of view. For this we need to study natural selection as a bottom-up forward process, using a mechanistic model that explains not only the natural selection of metabolic scaling but also the selection of metabolism and mass. Not only that, but to be sure that the overall selection is self-sustained, the explanation should not depend on other evolved traits that are part of the model but not explained by it.

Malthusian relativity (Witting 1995, 2008, 2017a,b) explains the natural selection of metabolic scaling based on this extended principle, where both the independent and dependent traits are predicted by a single integrated model of natural selection. As the independent traits of the dependent traits may depend on the natural selection of a deeper layer of independent traits, this stronger theoretical principle cascades into a paradigm of inevitable evolution by deterministic natural selection (Witting 1997, 2008). Given an abiotic environment suitable for life, inevitable evolution refers to the subset of biological evolution that follows from the natural selection that unfolds mechanistically from the origin of replicating molecules.

Malthusian relativity is based on this bottom-up forwardly unfolding natural selection, as it evolves from a selected increase in the net energy that is allocated to replication, generating a gradual unfolding of a population dynamic feedback selection by density dependent interactive competition. This selection involves a complete population ecological life history model, where the net energy that drives the feedback selection is obtained by an ecological foraging that optimises the trade-off between the cost of interference and the cost of local resource exploitation. This means that the explained allometric scaling involves other traits than just metabolism, including the life history and foraging ecology as a whole (Witting 1995, 2017a).

In a recent study I integrated the allometric component of Malthusian relativity with more than 25,000 estimates of the life history and ecological traits of mammals, estimating population ecological life history models for 4,936 species of mammals (Witting 2024). In the current paper, I use the trait variation of these models to illustrate how the majority of the inter-specific variation in the body masses, life histories, and ecological traits of mammals are reconciled by the interspecific variation in a few traits at the core of population dynamic feed-back selection. This explanation of the inter-specific variation involves allometric scaling, but it depends also on other factors like ecological variation in mortality and a selection attractor of interactive competition.

### 1.1 Bottom-up unfolding natural selection

In my analysis I decompose the inter-specific life history variation from a few primary drivers of population dynamic feedback selection, and this section identifies these main drivers.

Although most life history traits show a strong interspecific correlation with mass, body mass is not the primary driver of natural selection because its selection depends on other traits. Body mass is part of the quality-quantity trade-off (Smith and Fretwell 1974; Stearns 1992)—where a given amount of energy can produce a few large or many small offspring—and this selects for a continued decline in mass, when other things are equal.

The selection of mass in multicellular organisms is therefore dependent on an interactive competition where mass is selected as a competitive trait that is used by the larger than average individuals to monopolise resources. This occurs by a density-frequency-dependent selection, where the level of interference competition needs to be sufficiently high before the interactive selection of mass is stronger than the quality-quantity trade-off selection against mass.

The level of interactive competition that is required for this selection depends on an abundance that is so large that individuals meet sufficiently often in interactive competition, and this abundance depends on population growth with a quality-quantity balance that produces sufficiently many offspring from the net energy that is allocated to reproduction. The result is a population dynamic feedback attractor that selects mass in proportion to net energy by maintaining the invariant level of interference competition that is needed to balance the selection of the quality-quantity trade-off (Witting 1997; with the selection attractor of invariant interference called a competitive interaction fixpoint).

This feedback selection indicates that the net energy that is used on replication could be a primary driver of natural selection. Net energy, however, is not a completely independent trait, because a product between resource handling and the pace of handling defines it (Witting 2017a,b; with population dynamic feedback selection selecting the pace of handling as the pace of metabolism, in proportion to mass-specific metabolism).

A secondary mass-rescaling selection that occurs during the feedback selection of mass is another essential factor to consider in relation to the hierarchy of natural selection causalities (Witting 2017a). A potential selection increase in mass implies that the larger offspring metabolises more energy during the period of parental care, and variants that avoid this extra metabolic cost of the extra mass will be selected over variants that do not. This selects variants that reduce the metabolic need by a decline in mass-specific metabolism, generating the observed allometric downscaling of mass-specific metabolism with mass.

The available net energy per unit physical time, however, declines with a decline in mass-specific metabolism and this reduces the reproductive rate. This aggregated problem of selecting mass with mass-rescaled metabolism is solved by variants that dilate biological periods and thereby maintains the net energy and reproduction of the organism on the per-generation timescale of natural selection. This generates the observed inverse allometric scaling between periods/ages and mass-specific metabolism.

The numerical response of this mass-rescaling selection is captured by the exponents of the body mass allometries. These are selected by the optimal foraging in overlapping home ranges that generates the net energy for the overall feedback selection of mass, extending allometric scaling to ecological traits like home range and abundance. The result is a joint allometric scaling—of the metabolism, life history, and population ecology—that evolves as a sub-component of the natural selection of mass, instead of being a response to a physiological adaptation to size. For the original mathematical deductions of the allometric exponents see Witting (1995), for an extended deduction with primary selected mass-specific metabolism see Witting (2017a), and for a graphical deduction see Fig. 1.

**Figure 1:**
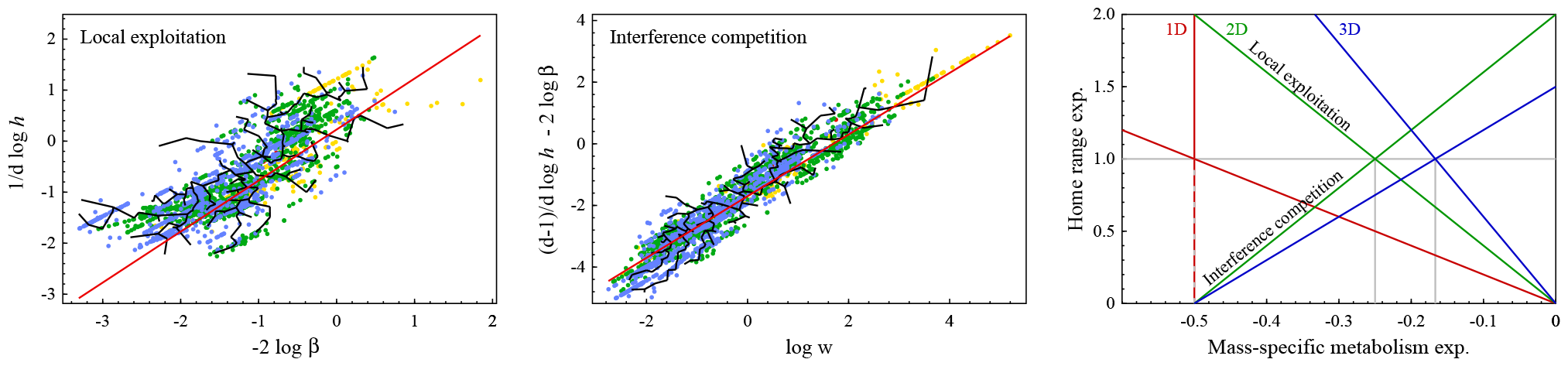
Allometric deduction. The left and middle plots show allometric representations of invariant regulations by local exploitation and interference competition, with black lines connecting data for placentals, coloured dots being placental estimates by estimators at different taxonomic levels, and red lines being the theoretical predictions. These predictions are solved in the right plot for the mass-rescaling exponents *ĥ* and 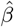 for home range and mass-specific metabolism. With *ĥ* increasing with 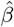 for interference competition 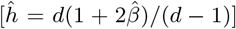 and declining for local exploitation 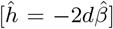, the selected exponents 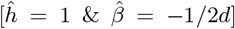 are the solutions where the lines with similar ecological dimensionality (*d* ∈ *{*1, 2, 3*}*) cross. From Witting (2023).

Resource handling is one of the few life history traits that are unaffected by mass-rescaling selection (Witting 2017a). This makes it evolutionarily independent of mass, with a primary selection that drives the evolution of other traits by its contribution to the net energy driven population dynamic feedback selection. Inter-specific variation in resource handling should thus explain large amounts of the variation in net energy and body mass, and secondarily also of the variation in other traits by their mass-rescaling dependence on the explained variation in mass.

Having removed the variance components that follow from variation in primary selected resource handling, I turn to the influence that the residual variation in the survival of offspring and adults have on the remaining life history variation. Ecological variation in mortality selects additional life history variation by perturbations of the competitive interaction fixpoint. An increase in mortality generates a decline in abundance and interactive competition, generating selection for increased replication until the interactive competition of the competitive interaction fixpoint is re-established. The energy for the selected increase in replication is taken primarily from body mass, with associated mass-rescaling selection for a wider range of life history variation.

Mass-specific metabolism is another potential life history influencer, as it is selected not only by secondary mass-rescaling but also by the primary selection that generates net energy for self-replication. The latter contributes to the feed-back selection of mass with superimposed mass-rescaling that downscales— at least to some degree—the primary selected mass-specific metabolism (Witting 2017a).

The importance of primary selected mass-specific metabolism for the selection of net energy and body mass is reflected in the values of the selected body mass allometries (Witting 2017a). Yet, mammals have approximate Kleiber (1932) scaling with typical 1/4-like exponents, and this agrees with a theoretical prediction where it is the variation in resource handling (and not metabolic pace) that generates the variation in the naturally selected body masses. This seems to be the case for the majority of multicellular animals, while the allometric scaling of unicellular eucaryotes and especially prokaryotes indicates a major influence from primary selected metabolism in these taxa (Witting 2017a,b). Yet, to check for a potential influence from primary selected metabolism also in mammals, following the decomposition from resource handling and ecological variation in mortality, I examine for a residual influence from mass-specific metabolism.

Having accounted for the selection influence from variation in net energy and mortality, I turn to the influence that the residual variation in the selection attractor of interactive competition has on the residual variation in other traits. With the most invariant component of the attractor being the intra-population fitness gradient in the cost of interactive competition, we expect some variation in our measure of interactive competition. Hence, the ecological traits that determine the level of interference (like abundance and home range overlap) should be selected to match the selected interference.

## 2 Methods

Following the selection hypothesis above, I decompose the inter-specific life history and ecological variation that Witting (2024) estimated for 4,936 species of mammals, covering the parameters in Table 1. This variation was estimated from 26,018 published trait estimates, with the inter-specific extrapolations of missing parameters following from the allometric correlations of the data.

**Table 1:** Traits. The analysed traits, their symbols (S), units, and order of magnitude (M) inter-specific variation with medians, 5th, and 95th quantiles. Estimates from Witting (2024).

| Trait | S | Unit | M | 5th | Median | 95th |
| --- | --- | --- | --- | --- | --- | --- |
| Body mass | $w$ | kg | 7.9 | 0.0098 | 0.16 | 95 |
| Res. handling | $\alpha$ | J | 8.8 | 0.0014 | 0.046 | 31 |
| F. metabolism | $\beta$ | W/kg | 2.6 | 2.5 | 14 | 46 |
| B. metabolism | $\underline{\beta}$ | W/kg | 2.7 | 1.2 | 4.6 | 16 |
| Net energy | $\epsilon$ | W | 6.8 | 0.045 | 0.68 | 95 |
| Incubation | $t_p$ | y | 1.9 | 0.05 | 0.086 | 0.68 |
| Fledging | $t_j$ | y | 2.5 | 0.05 | 0.09 | 0.96 |
| Rep. maturity | $t_m$ | y | 2.5 | 0.21 | 0.66 | 4.5 |
| Rep. period | $t_r$ | y | 2.5 | 0.33 | 1 | 9 |
| Generation | $t_g$ | y | 2.6 | 0.39 | 1.5 | 12 |
| Lifespan | $t_l$ | y | 2.7 | 1.6 | 5.2 | 35 |
| Off. survival | $l_m$ | $/t_m$ | 1.7 | 0.2 | 0.34 | 0.48 |
| Ad. mortality | $q_{ad}$ | $/y$ | 1.9 | 0.068 | 0.6 | 0.87 |
| Rep. rate | $m$ | $/y$ | 2.6 | 0.66 | 6.2 | 21 |
| Lifetime Rep. | $R$ | $/t_r$ | 1.7 | 4.2 | 5.8 | 10 |
| Abundance | $N$ | $/\text{km}^2$ | 7.1 | 0.57 | 290 | 2,900 |
| Biomass | $b$ | $\text{kg}/\text{km}^2$ | 5 | 2.2 | 41 | 740 |
| Pop. ene. use | $\epsilon_n$ | $\text{W}/\text{km}^2$ | 4.8 | 27 | 690 | 7,000 |
| Home range | $h$ | $\text{km}^2$ | 9.7 | 0.00027 | 0.0054 | 21 |
| Ho. overlap | $h_o$ | - | 6.4 | 0.12 | 2 | 55 |
| Interference | $\iota$ | - | 4.9 | -0.36 | 2.5 | 4.5 |

Some of the relevant traits are not available as data, and they were therefore calculated from other traits based on the trait relations in the population ecological model (Witting 2024). The variance decompositions that involve such traits are therefore not always independent of other traits, and this is discussed in the result section when relevant.

### 2.1 Explaining variance

I aim to explain the inter-specific variation within and across the 27 orders of mammals. To calculate how variation in net energy (*α & β*), mortality (*q*_*ad*_ *& l*_*m*_), and interactive competition (*ι*) explain the inter-specific variation, I use double logarithmic relations 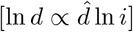 where exponents 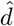 define the dependence of dependent traits *d* on the independent traits *i* ∈ {*α, q*_*ad*_, *l*_*m*_, *β, ι*}.

To predict a value 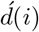, and calculate the associated residual value 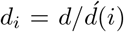, of a dependent trait *d* of a species in order *o*, I use a relation

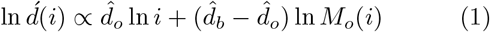

where *i* is the independent trait of the species, *M*_*o*_(*i*) is the median of *i* across the species in order *o*, 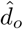 is the exponent that minimises the residual variance of ln 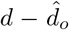 ln *i* across the species in the order, and 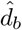 is the exponent that minimised the residual variance of 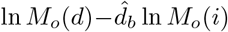 across the medians of the different orders. The within order exponents 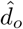 are estimated separately for all orders with life history estimates for more than *n* = 25 species, and set to the *n* weighted average of those within order exponents for orders with fewer estimates.

I present the values, and residual values, of traits as relative values (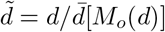 and 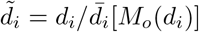) that are scaled by the average of the medians of the different orders, with residual values being calculated for the following sequence *α, q*_*ad*_, *l*_*m*_, *β*, and *ι* of the independent traits. The predictions are then evaluated by the average within-order variance of the dependent traits 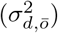 and their residuals 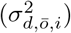, together with the between-order variance 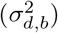, and residual variance 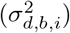, between the medians of the different orders.

To analyse the explained variance across traits, I use the proportion

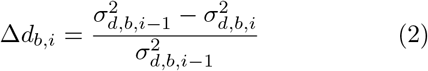

of the residual between-order variance in the dependent trait *d* that is explained by the independent trait *i*, and the corresponding proportion

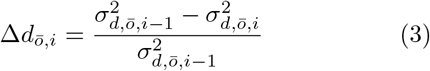

for the within-order variance. The total (*t*) proportion of the variance that is explained by all independent traits are 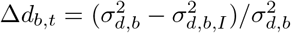 and 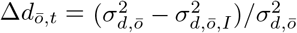.

To analyse for differences in the variance that is explained between and within orders, I use the proportional difference

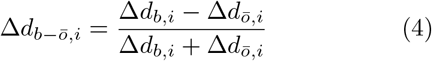

between the explained between and within order variance, with positive values implying that more variance is explained between orders than within, and negative values implying the opposite.

## 3 Results

Table 3 lists the estimated exponents (the average of the between order exponent and the *n* weighted average of the within order exponents) and the reduction in the within and across order variance of the different traits as a function of the independent trait components of *α, q*_*ad*_, *l*_*m*_, *β*, and *ι*. Fig. 2 illustrates these changes in the trait distributions of the different orders as the variance is explained by the independent traits (including for clarity only the cases where an independent trait explains 10% or more of the traits variance). The widths of the distributions reduce, and the medians of the different orders converge on the overall median, as the independent traits explain the variance.

**Table 2:**
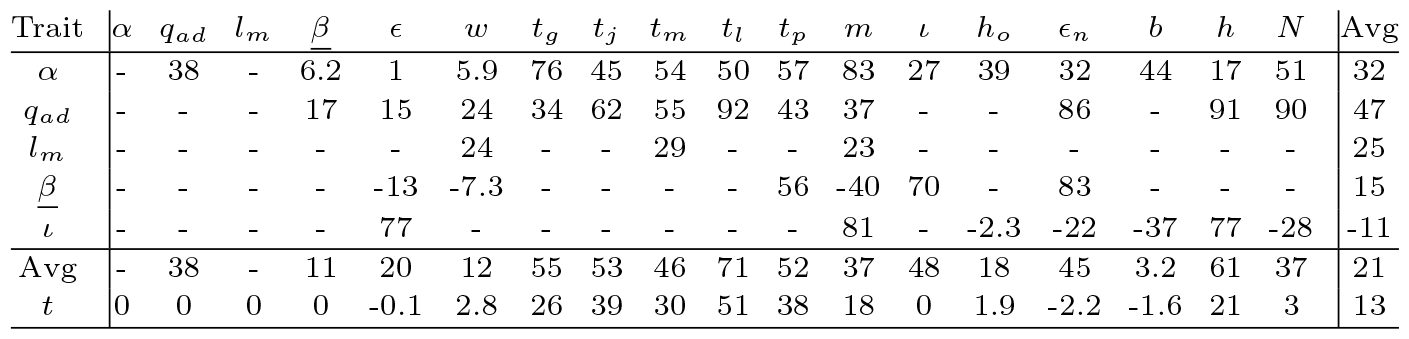
Between versus within order variance. The proportional difference (eqn 4, in percent) between the between and within order variance that is explained by the independent traits. Positive values are cases where more between order variance than within order variance is explained, and negative values are cases where more within than between order variance is explained. Calculated only for dependent traits where more than ten percent of the total variance is explained by an independent trait, with *t* denoting the joint effect of all independent traits *i* ∈ {*α, q*_*ad*_, *l*_*m*_, *β, ι*}.

**Table 3:**
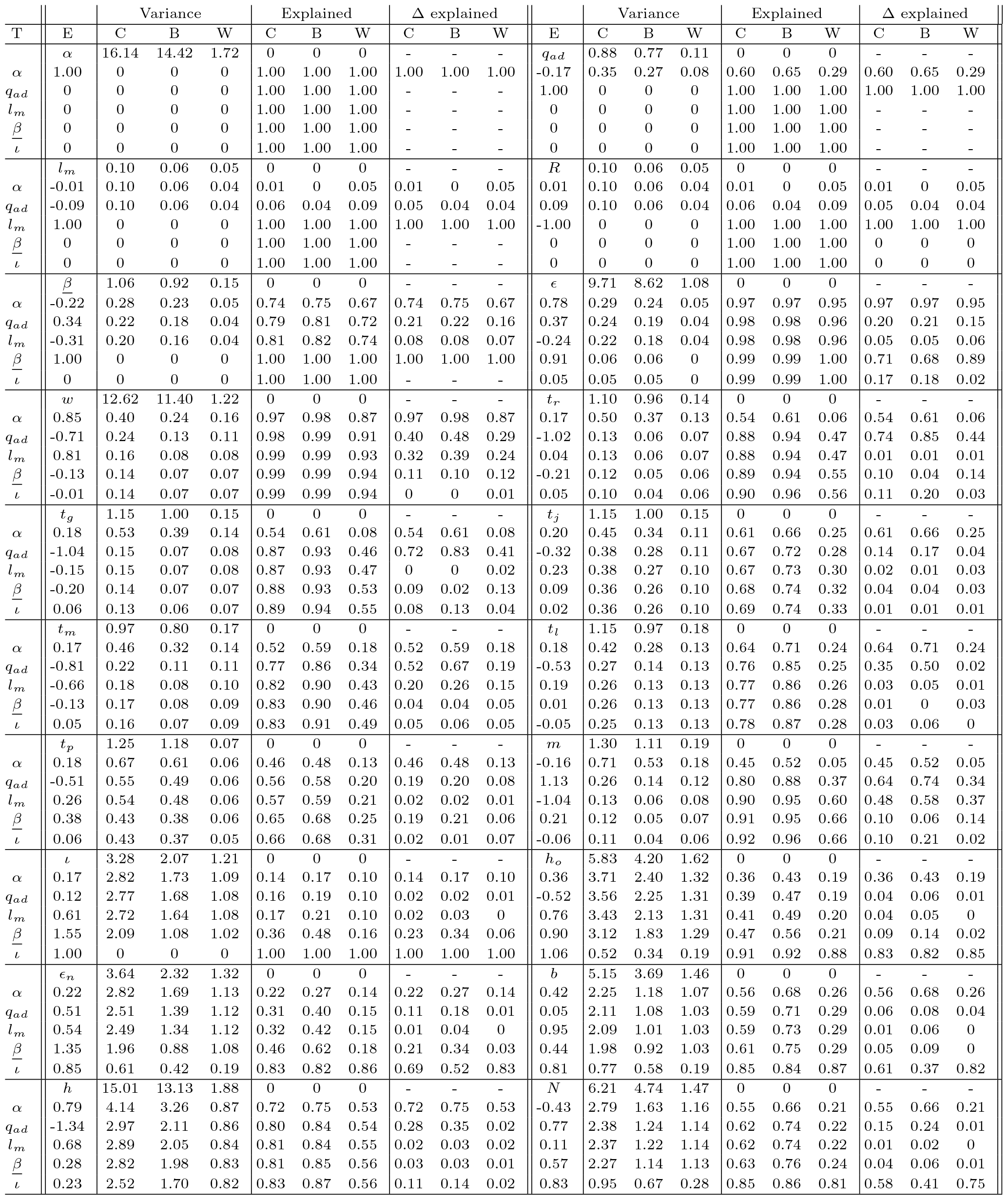
Explained variance. Variance: First rows: The between orden (B) variance, average within orden (W) variance, and combined variance (C=B+W) for the trait in columns ϵ. Other rows: The residual variance that is not explained by allometric correlations with the traits in column T. Explained: The fraction of the total variance that is explained. Δ explained: The fraction of the residual variance that is explained by the column T trait (eqns 2 and 3). The ϵ column values are the average of the between and within order exponents that explain most variance.

**Figure 2:**
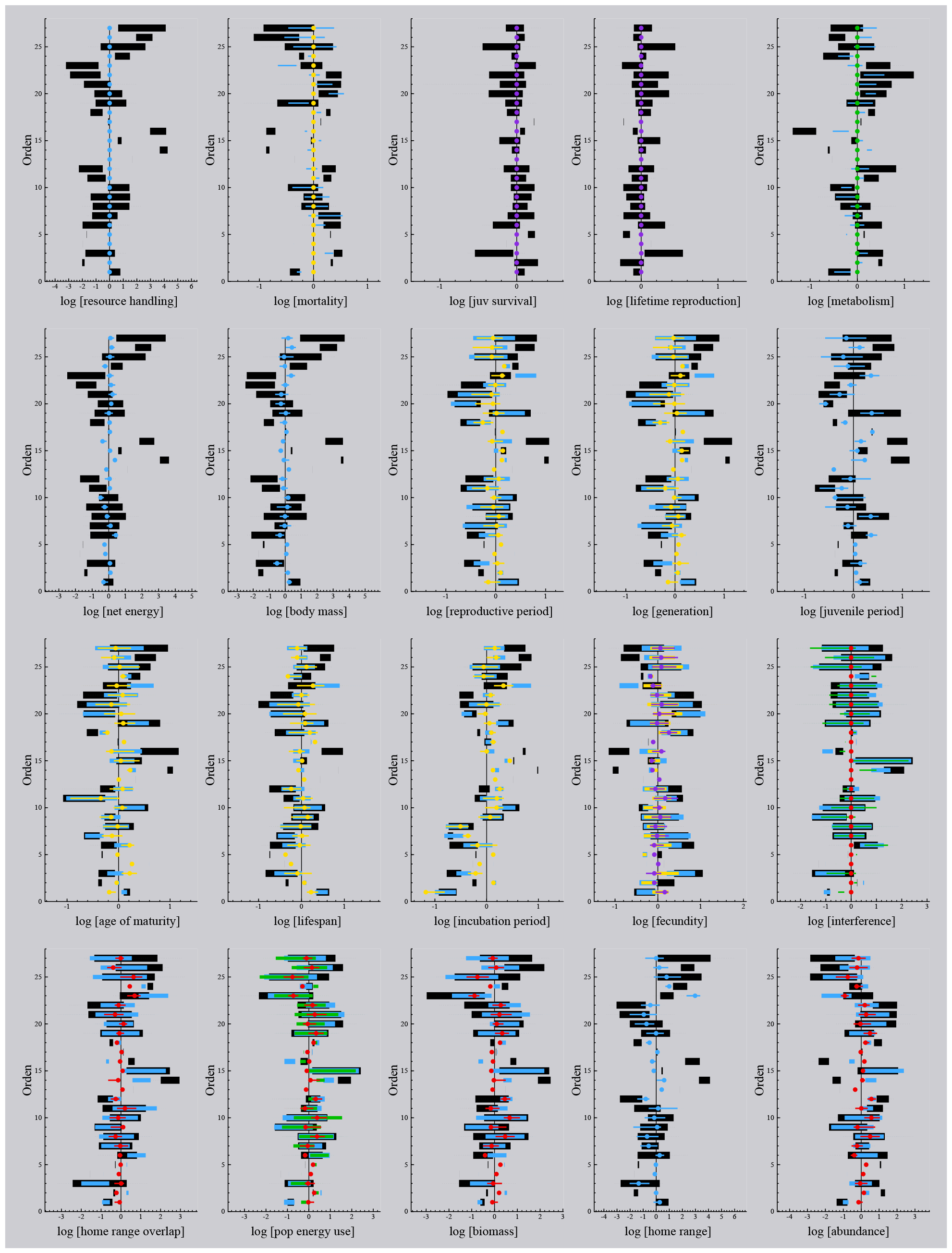
Explained variation. The 90% intervals (black bars), and limits (dashed lines), of the different traits across orders, including also the residual intervals and limits of the variation that is unexplained by resource handling (blue), residual adult mortality (yellow), residual juvenile survival (purple), residual metabolism (green), and residual interactive competition (red). Only cases where a dependent trait explains 10% or more of the variation are shown, with coloured dots being the medians of the residual variation for the last independent trait that explains more than 10%. Order: 1:Monotremata, 2:Paucituberculata, 3:Didelphimorphia, 4:Microbiotheria, 5:Notoryctemorphia, 6:Dasyuromorphia, 7:Peramelemorphia, 8:Diprotodontia, 9:Cingulata, 10:Pilosa, 11:Macroscelidea, 12:Afrosoricida, 13:Tubulidentata, 14:Proboscidea, 15:Hyracoidea, 16:Sirenia, 17:Dermoptera, 18:Scandentia, 19:Primates, 20:Lagomorpha, 21:Rodentia, 22:Eulipotyphla, 23:Chiroptera, 24:Pholidota, 25:Carnivora, 26:Perissodactyla, 27:Artiodactyla

My analysis reconciles variation between orders better than within-order variation (Table 2). The average deviations in the medians of orders that are explained by the independent traits is 91% compared to an average value of 55% for the within order explained variance.

### 3.1 Selection decomposition

#### Resource handling

Resource handling (*α*) generates net energy (*ϵ*) for the population dynamic feedback selection of mass, with the associated mass-rescaling selection inducing secondary effects on other traits (Witting 2017a). My estimates of resource handling, however, are estimated from net energy and mass-specific metabolism (*α* = *ϵ/β*), and net energy is estimated from the combustion energy of body mass (*w*_*ϵ*_), yearly reproduction (*m*), and metabolism (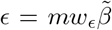, with 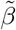 being a relative measure of the energy that is metabolised by offspring; Witting 2024). The variance decomposition from resource handling should thus not be seen as a statistical test, but only as a description of how the population ecological model reconciles the inter-specific life history variation with the different components of the net energy that generates the selection of the life histories.

For mammals, where the empirical allometries fit with body mass variation that is selected predominantly from variation in resource handling (Witting 2017a), net energy (*ϵ*) is predicted to scale as a ln *ϵ* ∝ 0.75 ln *α* function of resource handling, with body mass selected in proportion with resource handling (ln *w* ∝ 1.33 ln *ϵ* ∝ 1 ln *α*), and the second component of net energy (*ϵ* = *αβ*), i.e., mass-specific metabolism (*β*, metabolic pace), being a declining function of resource handling (ln *β* ∝ −0.25 ln *w* ∝ −0.25 ln *α*) owing to the selected mass-rescaling decline in mass-specific metabolism (ln *β* ∝ −0.25 ln *w*).

These expectations are reflected in the variance decomposition of the life history models, where resource handling raised to the 0.78 power accounts for 97% of the variance in net energy. 97% of the variance in body mass follows from a somewhat lower than proportional dependence on resource handling (0.85 exponent), and 74% of the variance in mass-specific metabolism is captured by resource handling raised to the −0.22 power.

Lower but large percentages are also explained for most of the remaining traits, including adult mortality (60%), generation time (54%), juvenile period (61%), reproductive maturity (52%), lifespan (64%), gestation period (46%), reproductive rate (45%), biomass (56%), home range (72%), and abundance (55%).

These explained percentages follow from the secondary effects of mass-rescaling. Given the 0.85 power dependence of body mass on resource handling, the expected and observed exponents are −0.25*\**0.85 = −0.21 and −0.17 for adult mortality, −0.25 *\** 0.85 = −0.21 and −0.16 for annual reproduction, 1 *\** 0.85 = 0.85 and 0.79 for home range, −0.75 *\** 0.85 = −0.64 and −0.43 for population abundance, and 0.25 *\** 0.85 = 0.21 and about 0.2 for most of the life history ages and periods. For the two traits that are expected to be invariant of body mass—i.e., for lifetime reproduction and the probability that an offspring survives to the reproductive age—we estimate average exponents around zero and only 1% variance explained.

#### Adult mortality

Variation in mortality affects the selection of several traits by a perturbation of the selection attractor of interactive competition (Witting 1997, 2008). Increased mortality selects for an increase in reproduction that compensates for the decline in abundance and interference competition that follows from increased mortality. This selection affects several other traits secondarily, as the net energy for increased reproduction is generated predominately from a selection decline in mass; with the associated mass-rescaling having potential effects on other traits.

This selection is reflected in the residual variation that is explained by the residual variation in adult mortality. 64% of the residual variation in annual reproduction, and 40% of the residual variation in mass, are explained by the residual variation in annual mortality, with the rate of reproduction increasing (estimated exponent of 1.13) and body mass declining (estimated exponent of –0.71) with increased mortality.

A cascading mass-rescaling effect—with higher metabolism and shorter life periods from the smaller masses of increased morality—was also found. The residual variation in adult mortality explains 21% (0.34 exponent) of the residual variation in mass-specfic metabolism. It explains 19% (−0.51 exponent) of the residual variation in the gestation period, 52% (−0.81 exponent) of the age of maturity, and 35% (−0.53 exponents) of the variation in lifespan (here you should not pay attention to *t*_*r*_ and *t*_*g*_ as these are partially calculated from *q*_*ad*_).

Smaller cascading mass-rescaling effects are also seen for the body mass dependent ecological traits, with 15% (0.77 exponent) of the residual variation in abundance, and 28% (−1.34 exponent) of the residual variation in home range, explained by the residual variation in adult mortality. Note, that the response of abundance is in the opposite direction of density regulation, where increased mortality leads to a decline in abundance. The observed increase follows population dynamic feedback selection, that compensates for the decline in abundance by the selection of net energy from mass to reproduction until the abundance and level of interactive competition of the evolutionary equilibrium is re-established. This involves the superimposed secondary mass-rescaling where the abundance increases from the increased mortality selected decline in mass.

#### Offspring survival

The probability (*l*_*m*_) that an off-spring will survive to the age of reproductive maturity is predicted to have a large impact on fecundity (*m*) and lifetime reproduction (*R* = *t*_*r*_*m*) through the survival versus reproduction compensation in population dynamic feedback selection. Yet, the potential effect *R* = 2*/l*_*m*_ is incorporated as a constraint in the estimated models given the assumption of stable populations, where *l*_*m*_*R* = 2 is the expected lifetime reproduction of a female.

The feedback selection compensation between survival and reproduction, however, operates through a reallocation of energy between reproduction and mass. As for adult survival, we find that increased offspring survival selects for an increase in body mass, with an estimated exponent of 0.81, and 32% of the residual variation in body mass explained by the residual variation in offspring survival.

With less than 10% of the residual variation explained for most of the remaining traits, a secondary mass-rescaling effect is not really detected. There is instead a negative relation (−0.66 exponent) between *l*_*m*_ and reproductive maturity, with 20% of the residual variation explained, most likely, from a survival probability that declines the longer offspring need to survive to reach the reproductive age.

#### Metabolism

Mass-specific metabolism affects rate-dependent traits, including the pace of resource handling that generates net energy for the selection of mass. With 81% of the variation in mass-specific metabolism being explained already by mass-rescaling from primary variation in net energy and mortality, there is some residual metabolic variation left to detect extra variation in net energy and mass. Yet, with 97% of the variation in net energy explained already by resource handling, we can expect only a very small influence on the life history from primary (i.e., non mass-rescaling selected) variation in mass-specific metabolism.

This is reflected in the variance decomposition, where mass-specific metabolism explains 71% of the residual variation in net energy, with the dependence being about proportional as expected (exponent of 0.91). Yet, while the dependence of residual variation in net energy on mass-specific metabolism is strong, primary selection on mass-specific metabolism accounts for no more than 1% of the total variation in net energy. The small but positive increase in net energy is not detected in body mass, where the exponent is negative (−0.13) with only 11% of the residual variation explained. There is also no consistent mass-rescaling effect from the residual metabolic variation.

#### Interactive competition

The level of intra-specific interference per individual (*ι*) is probably the most derived of all the traits considered, in the sense that the mechanistic generation of interference depends on a multitude of ecological and physiological traits that change with evolutionary modifications of the life history (approximated here by calculating interference as a function of abundance, home range, metabolism, and body mass). But, instead of being a derived trait that follows as a passive consequence of natural selection changes in other traits, the level of interference is one of the most central independent traits in population dynamic feedback selection. It is the overall selection attractor that controls the natural selection of the life history by selecting net assimilated energy between the demographic traits and mass, with the attractor itself being unaffected by the selected variation.

The attractor is referred to as the competitive interaction fixpoint, and it has a theoretical interference level of *ι*^*\*\**^ = 1*/ψ* when body mass in selected at an evolutionary equilibrium, and a theoretical level of *ι*^*\*s*^ = (4*d* − 1)*/ψ*(2*d* − 1) when mass is selected to increase exponentially at an evolutionary steady state [Witting, 1997; *ψ* is the intra-population gradient (around the average life history) in the cost of interference (e.g., different access to resources) per unit interference on log scale; subscripts *\*\** and *\*s* denote evolutionary equilibrium and steady state; *d* is the dominant spatial dimensionality of the foraging ecology]. The real selection invariant parameter is not the level of interference itself, but the intra-population gradient in the cost of interference [*ι*^*\*\**^*ψ* = 1 & *ι*^*\*s*^*ψ* = (4*d* − 1)/(2*d* − 1)]. This gradient—that favours the large and competitively superior individuals in the population—is selected to balance the fitness gradient of the quality-quantity trade-off.

With unconstrained selection being more likely to occur among the larger and ecologically dominant species, there might be some positive correlation where a larger fraction of the species that are situated at the *ι*^*\*s*^ attractor are those with the largest resource handling and/or metabolic pace. Yet, apart from this potential correlation, we expect a level of interference that is invariant with respect to the selected variation in the life history. This is also to a large degree what we find.

Even though the level of interference is calculated from abundance, home range, metabolism, and body mass, it is the most invariant traits, i.e., the trait that is least explained by the variance decomposition from net energy and mortality. Where 63, 81, 99, and 81% of the total variance in abundance, home range, body mass and metabolism have been explained so far by independent traits, only 36% of interactive competition is explained. It is evident that a large fraction of the explained variation in the sub-components cancel out, leaving the level of interference largely unaffected by the underlying variation in other traits.

Resource handling and mass-specific metabolism account for 14% and 23% of the variation in interactive competition, with a positive dependence (exponents of 0.17 and 1.55) as expected from the likely link between net energy and the likelihood of unconstrained selection. As expected from the selected invariance, the residual variation in the estimated level of interference explains on average only five percent of the residual variation in the demographic and physiological traits, including mass. Yet, with the variation in population density and home range overlap being selected to match the level of interference at the selection attractor, we find that the residual variation in interactive competition accounts for 58% of the residual variation in abundance and 83% for the overlap between home ranges, with 69% and 61% explained for the more derived traits of population energy use and biomass. Of these ecological factors, it is especially the home range overlap that relates most directly with the level of interference, as the probability to encounter other individuals is a direct function of the degree of overlap between home ranges. Somewhat surprisingly, only 11% of the residual variance in home range is reconciled by the variation in interactive competition. This may reflect that 81% of the variation in home range is already explained by the variation in net energy, survival, and metabolism, while this fraction is somewhat smaller for abundance (63%) and especially home range overlap (47%).

## 4 Discussion

Body mass is an essential evolutionary player that influences the evolution of other traits, but it is not the primary driver of natural selection as its selection depends on other traits. My analysis found variation in resource handling and mortality to reconcile 99% of the body mass variation in mammals. The associated secondary mass-rescaling explained 79% of the variation in metabolism, reproduction, and life periods/ages, and 72% of the variation in abundance and home range. All life history traits including body mass and metabolism, had no more than 1% of their variation explained by the level of interactive competition, which explained 83% and 58% of the residual variation in home range overlap and abundance. No consistent mass and mass-rescaling response could be detected from the residual variation in mass-specific metabolism, confirming that it is by far primary variation in resource handling that generates the body mass variation and allometric scaling in mammals.

The observed negative dependence of body mass on adult (−0.71 exponent) and offspring (−0.81 exponent) mortality is documented in several other studies, mainly for fish (Reznick et al. 1996; Haugen and Vøllestad 2001; Sinclair et al. 2002; Carlson et al. 2007; Herczeg et al. 2009). This supports the feedback selected allocation of net energy between reproduction and mass that maintains the naturally selected level of interference competition. This selection compensates for the decline in abundance that follows from a decline in population growth caused by increased mortality. The predicted, and observed, correlation is an increased abundance with increased mortality, reflecting the secondary mass-rescaling that follows from the selected decline in mass with increased mortality. Population dynamic feedback selection is a much better predictor than density regulation for the observed interspecific covariance between mortality and abundance.

Comparative methods and phylogenetic ecology (e.g. Promilsow and Harvey 1990; Harvey and Pagel 1991; Sibly and Brown 2007; Sibly et al. 2012; Dobson 2012; Brown et al. 2018; Burger et al. 2019) are often using a somewhat similar variance decomposition as applied in the current paper. Yet, these methods have no explicit focus on the underlying natural selection causality, as they have no bottom-up selection that shows how the independent traits evolve by a deeper more fundamental natural selection component. And life history differences by phylogenetic distance are the evolutionary outcome of natural selection and other processes of evolution, and not the cause of evolution (Reeve and Sherman 2001).

Where comparative methods capture the allometric importance of mass by non-causal correlation analyses, Rubner (1883) proposed metabolic scaling as the result of a physiological scaling to variation in size, with other life history and ecological allometries following as second order effects (Peters 1983; Calder 1984; Brown et al. 2004). But the physiological scaling hypothesis was never shown to be consistent with the natural selection of size, and nor necessary given the natural selection of size. By deducing the joint allometric scaling of the metabolism, life history, and population ecology from the optimal foraging behind the natural selection of size, Witting (1995, 1997) showed that physiological scaling hypotheses are unnecessary because the allometric scaling is selected already by the natural selection of mass.

These results were ignored when the physiological scaling hypothesis was relaunched (West et al. 1997) and praised (Gates and Gittleman 1997; Purvis and Harvey 1997; Williams 1997) with a new mechanism two years later. A rebuttal raised the above mentioned problems (Witting 1998), but this did not prevent a subsequent explosion of several new physiological scaling hypotheses with a narrow focus on metabolism (West et al. 1999a,b; Banavar et al. 1999; Dodds et al. 2001; Dreyer and Puzio 2001; Rau 2002; Fujiwara 2003; Demetrius 2003; Santillán 2003; Glazier 2010), and elaborate ecological side effects (Gillooly et al. 2002; Brown et al. 2004; Sibly et al. 2012; Humphries and McCann 2014).

Surprisingly none of these studies apparently questioned why they explained metabolic scaling from physiological adaptations and unexplained variation in mass, when metabolic scaling was explained already by the natural selection that explains the variation in mass (see Section 1.1 for mechanistic details). The underlying population dynamic feedback selection explains not only mass, metabolic scaling, and additional life history and ecological allometries (Witting 1995, 2017a), but it reconciles additional allometric issues (Witting 2023) including

*i*) 1/4 to 1/6 like transitions in the inter-specific exponents between terrestrial and pelagic species (Witting 1995, 2017a),
*ii*) a curvature where the inter-specific metabolic exponent of placental mammals increases from about 2/3 to more than 3/4 with an increase in mass (Kolokotrones et al. 2010; MacKay 2011; Witting 2018),
*iii*) a decline in the inter-specific exponent of metabolism from prokaryotes over protist and protozoa to multicellular animals (Makarieva et al. 2008; DeLong et al. 2010; Witting 2017a),
*iv*) a transition from −1/4 like inter-specific scaling of mass-specific metabolism in major animal taxa to invariant scaling across taxa (Makarieva et al. 2005, 2008; Kiørboe and Hirst 2014; Witting 2017b), and
*v*) a change from about 3/4 to 3/2 in the allometric exponent for the rate of body mass evolution, as documented by the fossil record covering 30 million to 3.5 billion years (Witting 2020).

While none of the physiological hypotheses are necessary for the existence of body mass allometries, they link physiological traits to metabolic scaling, allowing for different interpretations of the cause and effect of selection as this was never resolved by the contingent approach in the first place. This allows the top-down selection of physiological adaptation to meet the bottom-up population dynamic feed-back selection of allometric scaling, providing prospects for a unified theory where the physiology adapts with the selected metabolism, mass, and allometric scaling.

